# A Dipolar Photoswitch Modulates Bacterial Membrane Potential and Reveals Context-dependent Bioelectrical Circuitry

**DOI:** 10.64898/2026.01.20.700538

**Authors:** Pietro Bertolotti, Fabio Marangi, Edoardo Cianflone, Andrea Pianetti, Helena R. Keller, Arianna Magni, Valentino Romano, Marta Ghidoli, Tailise Carolina de Souza-Guerreiro, Chiara Bertarelli, Guglielmo Lanzani, Munehiro Asally, Giuseppe Maria Paternò

## Abstract

Dynamic bioelectric signalling in bacteria regulates physiology and collective behaviours, yet tools to perturb microbial membrane voltage with high spatiotemporal control remain limited. Here we introduce MTP2, a non-genetic, membrane-targeting azobenzene photoswitch that enables optical modulation of bacterial membrane potential by tuning interfacial electrostatics in *Bacillus subtilis*. MTP2 associates strongly with the cell envelope and shifts the resting potential to more negative values in the dark, while 470-nm illumination evokes a robust, reversible depolarization at the single-cell level. Although MTP2 photoisomerization is ultrafast (picoseconds), the voltage waveforms unfold over seconds to minutes, indicating that the response is set by homeostatic ion transport rather than by MTP2 photochemistry. Genetic and pharmacological perturbations show that K^+^ conductance, Cl^-^-sensitive pathways, and active transport reshape the amplitude, kinetics and even polarity of the optical response, revealing a context-dependent interplay between a passive molecular perturbation and endogenous bioelectric circuitry. As a functional proof of concept, kanamycin efficacy co-varies with the optically tuned voltage state. Together, these results establish MTP2 as a reversible chemical optostimulator for probing and controlling microbial electrophysiology.

## 1. Introduction

The ability to generate and propagate electrical signals has traditionally been considered a characteristic feature of specialized excitable eukaryotic cells, such as neurons, cardiomyocytes and muscle cells. However, recent discoveries have demonstrated that bacteria also exhibit an active bioelectric behaviour, exploiting ion channels and membrane potential dynamics to coordinate behaviour and regulate key physiological processes.^1–4^ Far from being passive organisms, bacteria employ bioelectric signals to manage community behaviours, respond to environmental cues, and maintain internal homeostasis. A growing body of evidence underscores bioelectricity as a fundamental motive force shaping the genetics, physics, and physiology of bacteria.^5^

Central to bacterial bioelectricity is the electrical membrane potential, which results from the regulated flux of ions across the plasma membrane.^6^ Recent studies revealed that the bacterial membrane potential can be modulated by electrical stimulation^7,8^ and its dynamics underlies phenomena such as long-range electrical signalling within bacterial communities,^9–15^ thus enabling coordinated metabolic oscillations and resource sharing among cells. In motile bacterial colonies, electrical signals also influence motility by modulating ion gradients that drive flagellar rotation.^13,14^ Moreover, bacterial responses to antibiotics are increasingly linked to membrane potential dynamics, highlighting a key role for bacterial bioelectricity in survival under stress.^8,16–19^ Therefore, the emerging picture is that bacterial ion channels function as integral parts of a bioelectric communication network, akin to primitive nervous systems.^20,21^ This excitation-based route gains relevance under fluctuating environments or antibiotic pressure, where swift, collective decision-making can determine whether a population survives or is eradicated.^22^ Despite this intriguing scenario, prokaryotic electrophysiology remains underexplored compared to its eukaryotic counterpart, due to the lack of established tools for modulating and interrogating bacterial bioelectric behaviours. This gap underscores the pressing need for innovative bioengineering strategies to control membrane potential in real time.

Light-based approaches have become powerful tools for probing and modulating bacterial bioelectric phenomena,^21,23^ offering advantages over some intrinsic limitations of electrode-based stimulation and interrogation techniques. In particular, it allows achieving high spatiotemporal resolution that is crucial for targeting these small, motile, and highly heterogeneous organisms. Materials-driven synthetic biology, through the use of small molecules or engineered nanomaterials, has provided several routes for non-genetic, optically controlled modulation of cellular electrophysiology.^24–29^ These strategies can contribute to understanding bacterial bioelectricity under physiologically relevant conditions, as well as holding potential for translational applications, such as targeted disruption of biofilms or enhancement of antibiotic therapies.^30^

In this work, we implement a chemical optostimulation strategy using MTP2, an amphiphilic membrane-targeting azobenzene photoswitch, to transiently perturb membrane electrostatics via dipole-driven charge redistribution. This approach is mechanistically distinct from our optomechanical photomodulator Ziapin2, which modulates voltage primarily through membrane thickness changes.^25,31–34^ In neurons and cardiomyocytes, MTP2 partitions into the plasma membrane and illumination elicits depolarization, consistent with photoisomerization-driven changes in the molecular dipole and the effective surface-charge balance.^35,36^ Here we ask whether analogous optical perturbations can tune bacterial membrane potential in *Bacillus subtilis*, and we dissect how endogenous K^+^, Cl^-^, and active-transport pathways buffer, reshape, and in some contexts invert the resulting voltage response.

We first show that MTP2 passively and strongly associates with *B. subtilis* and retains ultrafast trans–cis photoisomerization in the membrane environment. We then demonstrate that MTP2 shifts the resting membrane potential to more negative values in the dark, while 470 nm illumination elicits a robust depolarization of ∼10–12 mV at the single-cell level. Motivated by the fact that voltage shifts can arise from comparatively small net charge imbalances across the membrane capacitor, we interpret these responses within a pump–leak framework in which MTP2 acts as a photo-tunable bias to the effective membrane “circuit” while native conductances and transport processes set the operating point and timescales. Using genetics and pharmacology, we show that the resulting voltage dynamics depend on an interplay between YugO K^+^ channels, chloride-sensitive pathways and active transport, which together buffer, amplify or invert the MTP2-driven response depending on context. Finally, we show that kanamycin efficacy tracks the optically tuned membrane potential: more negative *_m_* states increase kanamycin impact, whereas light-driven depolarization partially attenuates it, dissecting a functional link between externally imposed voltage dynamics and an antibiotic response.

## 2. Results

### 2.1 Molecular Structure and Optoelectronic Properties of MTP2

MTP2 is classified as a push-pull azobenzene bearing a nitro-substituted phenyl ring (a strong electron-withdrawing group) at one end and an aniline (electron donor group) N,N-disubstituted with ω-pyridinium-terminated alkyl chain at the other. The cationic heads confer amphiphilic character to the photochrome. In addition, such units can interact electrostatically with phospholipid head groups of the membrane. This D–A configuration supports intramolecular charge transfer, resulting in significant molecular dipole formation and unique optoelectronic properties (**Fig. 1A**).^37^

**Figure 1.**
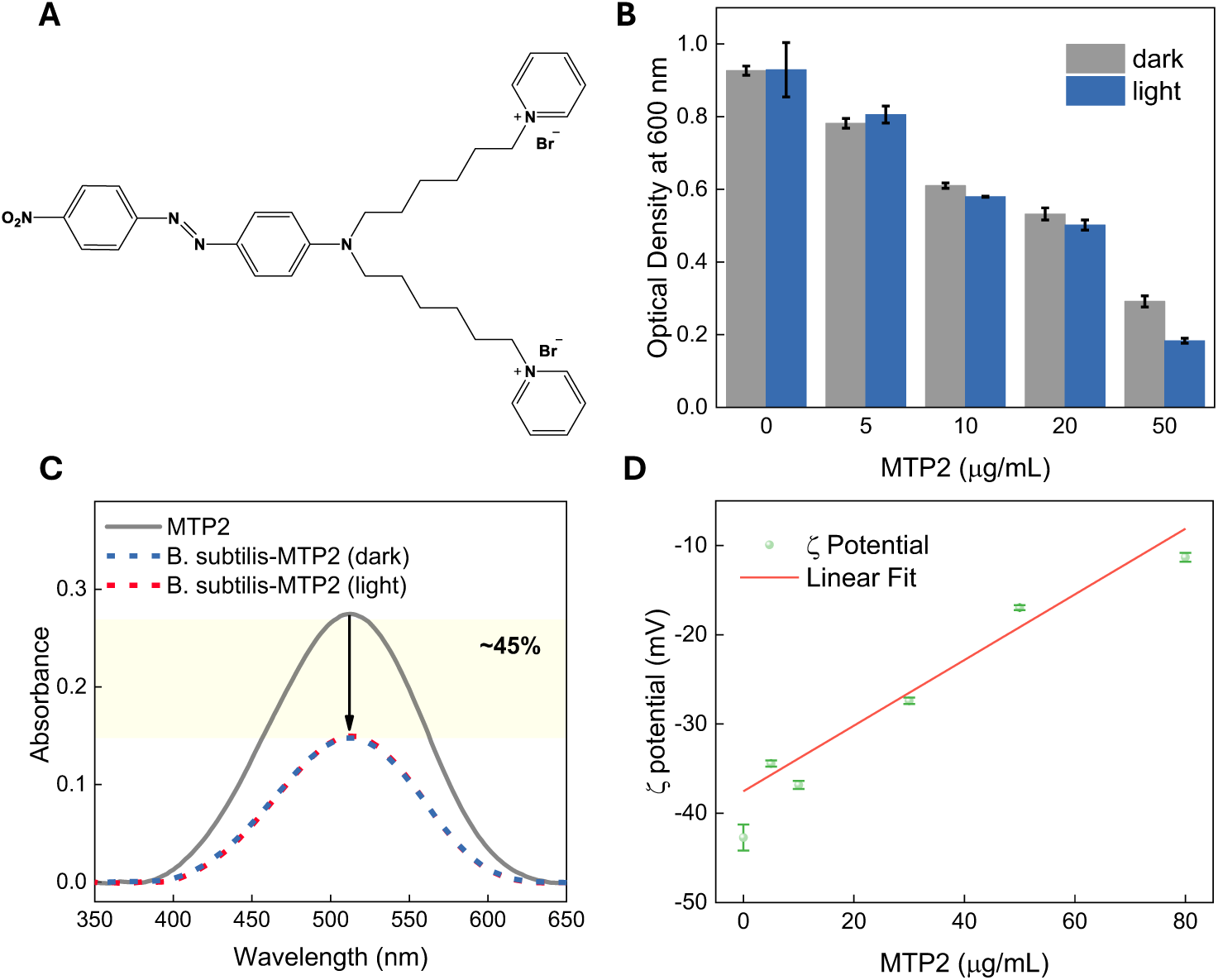
MTP2 uptake in B. subtilis and toxicity. (A) Molecular structure of MTP2. (B) Viability of B. subtilis exposed to MTP2 under dark and light conditions (470 nm), as measured by the final optical density at 600 nm. For these experiments, bacteria samples were exposed to λ = 470nm continuous wave light 10 min every 20 min of dark for 8 hours (power density = 11.5 mW/cm^2^). For every sample exposed to light, another one is kept in dark environment (control). The error bar represents the standard deviation over 6 different samples analysed in groups of 3 processed on 2 different days. (C) MTP2 uptake assay in B. subtilis cells at 10 µg·mL^−1^ in dark and under light conditions (power density = 11.5 mW/cm^2^, 10 min of illumination followed by 20 min in dark, repeated). (D) ζ -potential measurements of B. subtilis cells exposed to increasing concentrations of MTP2. The red line is linear fit of the data.

Under dark conditions, MTP2 adopts a *trans (E)*-configuration, with its molecular dipole moment oriented along the backbone connecting the donor and acceptor groups. Upon stimulation with 470 nm light, the molecule undergoes *trans-cis/(E-Z)* isomerization, which is accompanied by a significant decrease in the molecular dipole moment, approximately 30%.^35^ When the light stimulus is removed, MTP2 spontaneously reverts to its initial trans configuration over time, restoring the original D-A distance and dipole moment.^35^ In eukaryotes, membrane surface charge redistribution results from the overall re-orientation of the effective dipole deriving from the molecular ensemble, driving membrane potential depolarization.

### 2.2 Viability, uptake and surface association assays

We first assessed MTP2 toxicity under both light and dark conditions. *B. subtilis* cultures (OD_600_=0.1) were treated with increasing concentrations of MTP2 (0–50 µg· mL^−1^). Half of the cultures were exposed to intermittent 470 nm illumination (cycles of 10 min light, 20 min dark) during 8 h of growth. MTP2 induced a clear concentration-dependent toxicity, with significant inhibition of bacterial growth observed above ∼10 µg· mL^−1^ based on final optical densities (**Figure 1B**). Under these conditions, 470 nm illumination on cells loaded with MTP2 did not measurably affect viability. On this basis, 2–10 µg· mL^−1^ MTP2 was selected for subsequent photomodulation experiments.

We next tested whether MTP2 associates with the bacterial envelope/membrane using two complementary assays. First, we quantified uptake/retention by UV–Vis spectroscopy (**Figure 1C**). After incubating cells with 10 µg· mL^−1^ MTP2 for 1 hr at 25°C and pelleting,^38^ the absorption spectrum of the supernatant exhibited a marked decrease relative to the initial solution, indicating that a substantial fraction of MTP2 is retained by the cells. This behaviour is consistent with passive yet strong association of the di-cationic amphiphile with the Gram-positive cell envelope, and we did not observe evidence for a light-dependent change in uptake under the conditions tested. Using the retention fraction from the UV–Vis assay (e.g., ∼45% of 10 µg mL^−1^ retained) and a standard OD-to-cell conversion for *B. subtilis* (OD₆₀₀ ≈ 1 corresponds to ∼ 10^8^ – 5×10^8^ cells mL^−1^), this corresponds to an order-of-magnitude ∼ 10^6^ – 10^7^ molecules per cell (see SI for assumptions and calculation). By comparison, parent membrane-targeting azobenzenes have been reported to load eukaryotic cells at ∼ 10^3^–10^5^ molecules per cell.^39,40^

Second, ζ-potential measurements revealed a dose-dependent shift in the electrokinetic surface potential of cell suspensions upon MTP2 treatment (**Figure 1D**): increasing MTP2 concentrations made the ζ potential progressively less negative. This trend supports a dominant electrostatic component to MTP2–envelope association, consistent with partial screening/neutralization of the highly anionic surface of *B. subtilis*.^41^ Collectively, UV–Vis retention and ζ-potential shifts indicate that MTP2 pre-concentrates at the Gram-positive envelope through a dominant electrostatic component. The key implication is not only that MTP2 is cell-associated, but that its effective surface density at the membrane interface can be substantially enhanced relative to the bulk concentration, providing a quantitative basis for the larger voltage responses observed in bacteria than in eukaryotes (*vide infra*).

### 2.3 Optical spectroscopy characterization of MTP2

As MTP2 has a very fast isomerization, to characterize the photoisomerization of MTP2 under physiologically relevant conditions we performed sub-nanosecond transient absorption (TA) spectroscopy on MTP2 both in phosphate-buffered saline (PBS) and directly on suspensions of MTP2-treated *B. subtilis* cells in PBS. In TA, the dynamics of photoexcited molecules are monitored through a two-pulse scheme. First, an ultrafast “pump” pulse deposits energy into the sample, promoting a fraction of molecules from the ground state into electronically excited states. After, at a certain delay, a weaker white “probe” pulse interrogates the sample’s absorption spectrum. By systematically sweeping the pump–probe delay, a time-resolved map of how the excited-state populations evolve was constructed. In the resulting TA spectra, positive ΔT/T signals arise when the probe experiences increased transmission: either because the ground state has been depopulated (ground state bleach, GSB) or because excited molecules emit photons that add to the probe beam (stimulated emission, SE). By contrast, negative ΔT/T signals indicate decreased transmission, attributable to photoinduced absorptions (PIA), which correspond to additional absorption from the excited states. Together, these features reveal the lifetimes and interconversion kinetics of transient species generated by the pump pulse.

In both PBS and in PBS with *B. subtilis* (**Figure 2A,B**), the MTP2 TA spectra display a positive ΔT/T feature centred at ∼480 nm, coincident with the steady-state absorption band of the trans isomer and therefore assigned to ground-state bleach (GSB), as well as photoinduced absorption (PIA) band lying at 425 nm that can be attributed to the absorption of the cis isomer. In addition, we can observe a broad PIA band red-shifted with the respected to the GSB signal that can be assigned to absorption processes from the excited states. Kinetic traces of the GSB at 480 nm (**Figure 2C**) are well described by a fast decay followed by a long-lived residual signal. The fast component has time constants τ_1_ ≈ 1.8 ps in PBS and τ_1_ ≈ 2.3 ps in B. subtilis and reflects the decay of the initially excited trans population, which can either undergo trans → cis isomerization or relax back to the trans ground state without isomerizing. The subset of molecules that reach the cis ground state generates a depleted trans population, observed as the long-lived GSB.

**Figure 2.**
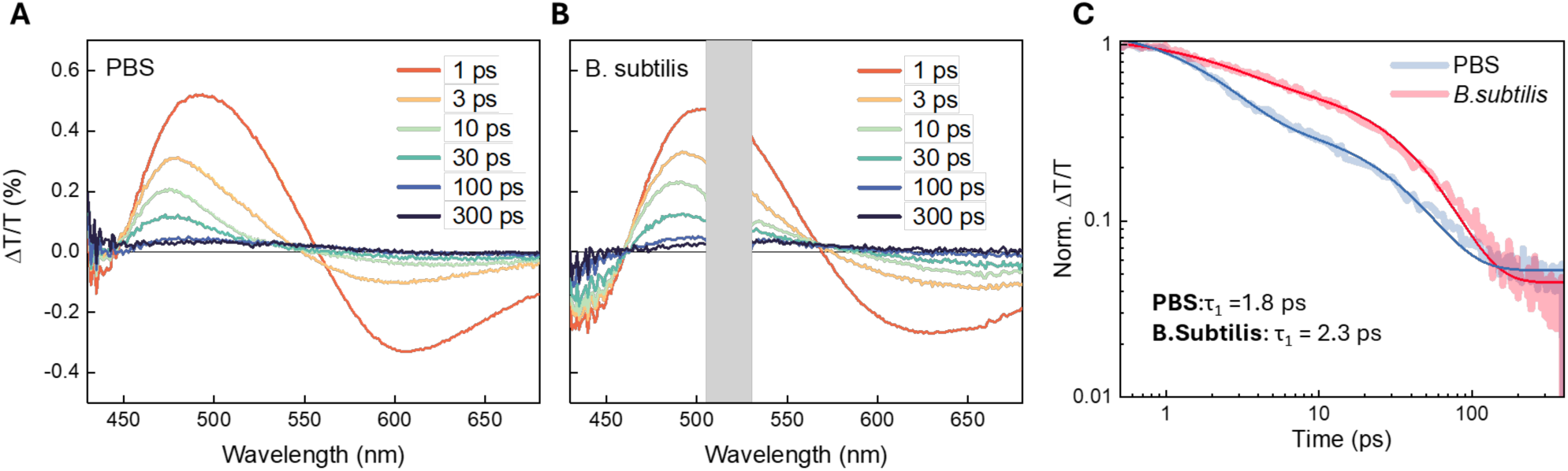
MTP2 Ultrafast Transient Absorption in B. subtilis. (A) TA spectra at different pump-probe delays for MTP2 in PBS and (B) in B. subtilis cells suspended in PBS to OD_600nm_ = 0.5. (C) Dynamical traces at a probe wavelength of 480 nm corresponding to the GSB region for MTP2 in PBS and B. subtilis cells. The excitation wavelength is 520 nm for both samples. Experimental data are reported as faded lines, while solid lines represent fittings with exponential functions. The grey rectangle in B is meant to hide the laser-driven scattering effects in the TA spectra of bacteria.

By comparing the amplitude of this long-lived bleach at late delays to the initial GSB amplitude, we estimate that ≈ 5.0 % of MTP2 molecules reside in the cis photoproduct state at the nanosecond scale in PBS, and ≈ 4.5 % in the *B. subtilis* suspension. These values correspond to the fraction of molecules that remain in the photoproduct state at this timescale; any cis isomer that decays back to trans faster than a few hundred picoseconds would not contribute and would make our estimates a lower bound to the initial isomerization yield. However, previous sub-ns–ms TA experiments on MTP2 in aqueous and membrane-mimicking environments have shown that the cis isomer relaxes on the tens-to-hundreds of microseconds timescale, far slower than our current window. We therefore take the residual bleach as a good figure of merit for the cis yield.^35^ Under this assumption, a comparable fraction of MTP2 molecules undergoes trans → cis isomerization in buffer and in the bacterial environment, with only a modest (∼10 % relative) reduction of the cis yield in cells.

### 2.4 MTP2 drives membrane potential dynamics in *B. subtilis* wild type

Following confirmation of MTP2’s membrane interaction and concentration-dependent lethality, its light-mediated membrane potential modulation was investigated by epifluorescence time-lapse microscopy with tetramethylrhodamine methyl ester (TMRM) as a voltage sensitive probe.^1,2,42–44^ Cells were exposed to MTP2 at concentrations of 2, 5, and 10 µg·mL^-1^.

In dark conditions, a progressive hyperpolarization was observed as the MTP2 concentration increased, with the average membrane potential shifting from −70 mV to −95 mV between 0 and 10 µg·mL^−1^ (**Figure 3A,B**). In parallel, we have seen that the ζ-potential of *B. subtilis* became less negative upon MTP2 addition, consistent with the incorporation of the di-cationic photoswitch at the cell envelope. This behaviour can be rationalized by a simple electrostatic picture. The positively charged headgroups of MTP2 partially neutralize the negative surface charge, rendering the outer surface potential (ζ) less negative, while the oriented intramembrane dipole of MTP2 increases the electric field across the bilayer and makes the cytosolic side more negative. In other words, the same population of embedded dipoles simultaneously decreases the apparent surface charge and increases the transmembrane potential difference, leading to the observed dark-state hyperpolarization (**Figure 3C**).

**Figure 3.**
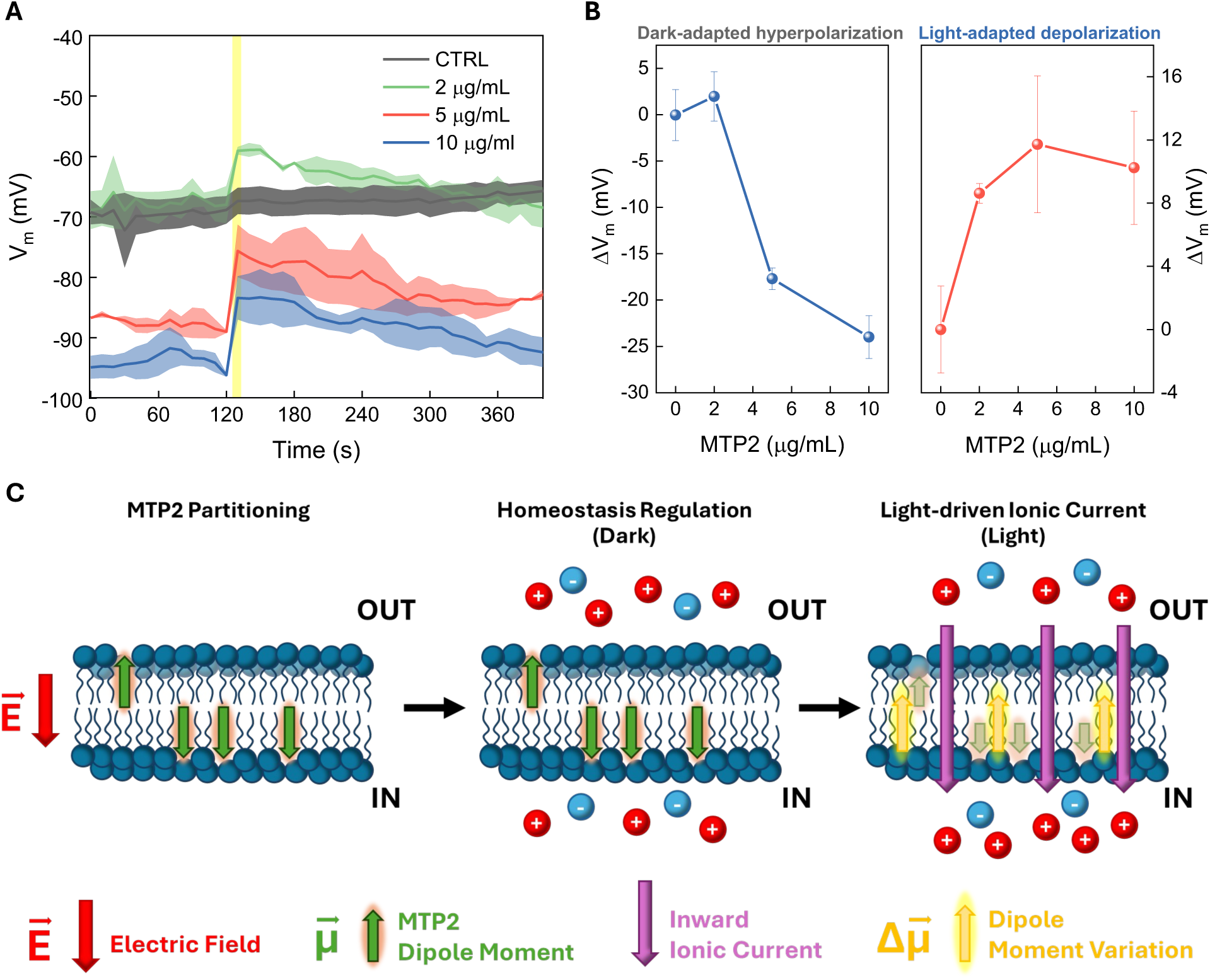
MTP2 effect on B. subtilis Electrophysiology. (A) Electrophysiological traces of B. subtilis upon administration of increasing concentrations of MTP2, before and after (yellow vertical line = 10 s) transient illumination at 470 nm. The halo represents the standard deviation (B) Membrane potential variation in B. subtilis at increasing MTP2 concentrations under dark conditions (left) and illumination (right), with error bars representing the standard deviation over 9 samples per condition processed on three different days and from different mother cultures. The ΔV is calculated from the difference between the MTP2 treated bacteria and the untreated ones. (C) Schematic working model for MTP2-driven voltage dynamics in B. subtilis. The molecule experiences an asymmetric electrostatic bias across the membrane (transmembrane field and interfacial surface potential), which can favour a preferred orientation/partitioning (left). In the dark, this perturbation acts as a steady bias to the membrane “circuit,” shifting the pump–leak balance toward a more negative operating point (hyperpolarized V_m_; centre). Upon transient 470 nm light exposure the MTP2 dipole moment 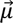 decreases yielding to a 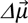 (right), transiently changing the interfacial bias and driving a net inward compensatory current that manifests as depolarization (right)

Intriguingly, a comparable dark hyperpolarization was not apparent in mammalian cells.^35^ This difference does not necessarily imply a different molecular functioning mechanism in the two cell types, but rather different ‘bioelectric gain’. In a capacitor picture, Δ*V_m_* = Δ*Q*/*C* = Δ*σ*/*C_m_*, and *C_m_* (specific capacitance per unit area) is of the same order for most lipid membranes.^2^ Thus, the amplitude of the response is expected to depend primarily on (i) the net interfacial perturbation per area (Δσ), which can be increased by electrostatic pre-concentration of MTP2 at the highly anionic Gram-positive envelope, and (ii) the effective pump–leak buffering that converts a given bias current into a sustained Δ*V_m_* (see S.I. for calculations and assumptions).

Upon a 10-second pulse of 470 nm light, cells exhibited a measurable depolarization peak of approximately 12 mV (**Figure 3A, B**), occurring within 10 seconds of light exposure. This finding is consistent with the proposed mechanism, whereby illumination reduces the dipole moment, which diminishes hyperpolarization and produces depolarization (**Figure 3C**). An interesting aspect was the slow recovery of the membrane potential to the hyperpolarized resting state after the light-induced depolarization, which occurred on the scale of minutes, similar to responses observed previously with Ziapin2.^25^ This slow metabolic adjustment is not fully attributed to the specific nature of the azobenzene stimulus (ensemble re-equilibration occurring in ∼ hundreds of µs) or TMRM equilibration time (∼30 s),^42^ but rather to an overall slower bioelectric machinery in bacteria (minutes) compared to, for instance, neurons (milliseconds).^31^ In summary, MTP2 enables biphasic modulation of membrane potential: in the dark the trans dipole induces hyperpolarization in a MTP2 concentration-dependent manner; under light the dipole change triggers depolarization. This demonstrates that MTP2 photoswitching can non-invasively control *B. subtilis* membrane potential in single cells.

### 2.5 Role of Potassium Ion Channels (YugO channels)

In mammalian cells, the effect of MTP2 was primarily attributed to the modulation of surface charge due to dipole moment variation-induced charge distribution and ion flux. However, in *B. subtilis*, we aim to investigate whether the ion transport mechanisms, driven by the bacterial bioelectric machinery, are involved in the observed responses. Since potassium flux is a primary determinant of bacterial membrane potential, we examined its role using a *B. subtilis* mutant (*ΔyugO*), lacking the gene encoding for the K^+^ channel YugO. ^9,11^ This channel plays a crucial role in biofilm formation and electrical communication though K^+^ efflux.^9,45^ It is worth noting that a concentration of 5 μg/mL MTP2 was used in these experiments, a level at which both dark- and light-adapted electrophysiological effects begin to show significant changes.

The dark-state hyperpolarization (**Figure 4A**) was similar to that of wild type (WT) exposed to MTP2 (Fig.3A), indicating that the YugO channel is only marginally involved in maintaining the basal membrane potential balance in the presence of MTP2 at steady state. On the other hand, light-induced depolarization was markedly enhanced in *ΔyugO* strain (ΔV ∼ +30 mV peak *vs*. +12 mV in WT), suggesting that YugO channel contributes significantly to buffering or limiting membrane depolarization during transient stimulation. When cells experience a light-induced depolarization due to MTP2 dipole moment variation, the opening of YugO channels would facilitate potassium flux, thereby restraining excessive depolarization. In *ΔyugO*, this compensatory mechanism is lost due to the absence of these K^+^ channels, resulting in an exaggerated, unregulated light-triggered depolarization.

**Figure 4.**
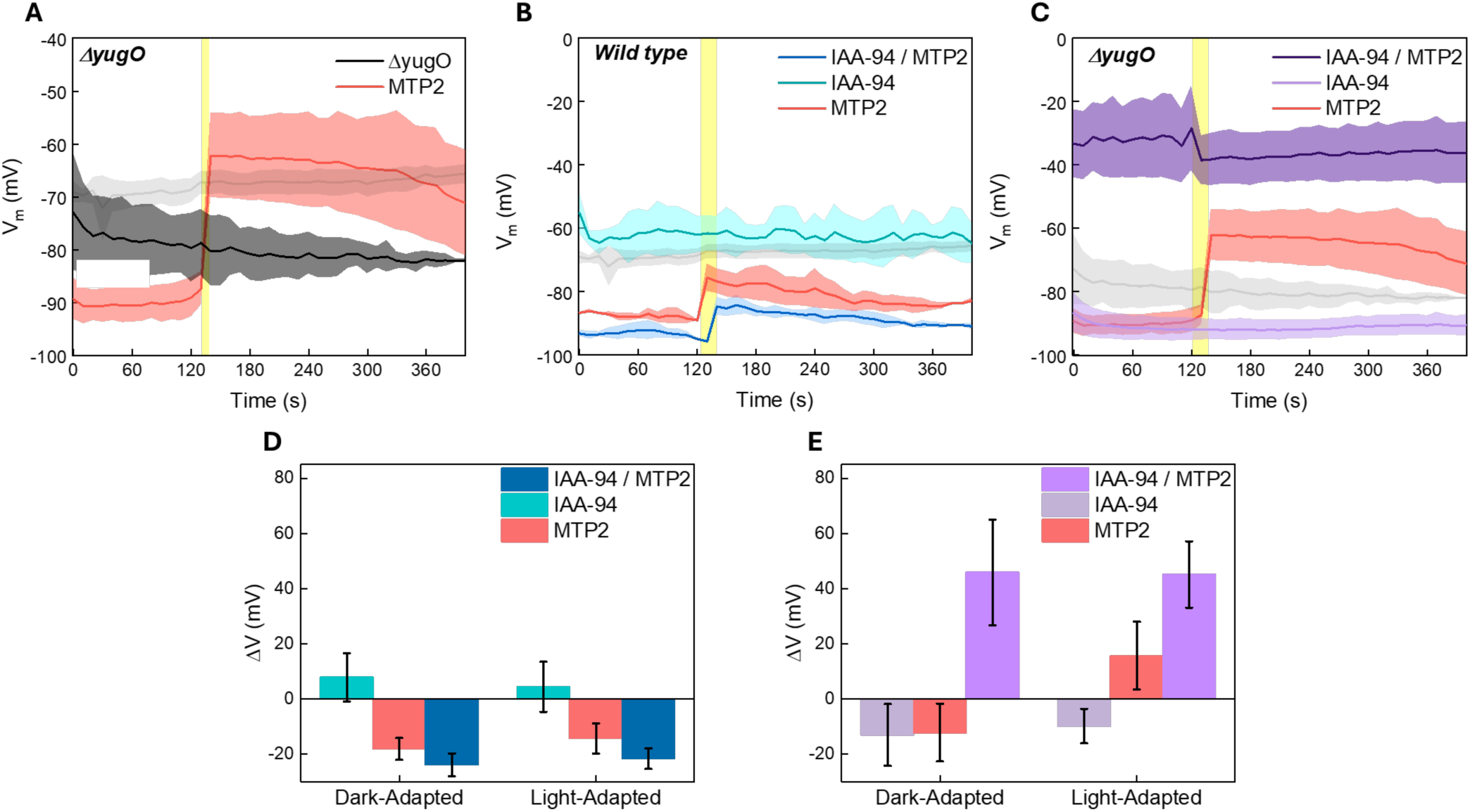
MTP2 affects passive K^+^ transport and chloride channel blockers modulatory role. Electrophysiological traces of B. subtilis lacking the YugO channel (ΔyugO) upon administration of MTP2 at 5 μg/mL (A). Light onset is represented by a yellow vertical line (10 s). The halo represents the standard deviation. The untreated WT condition is reported in transparency (grey). Electrophysiological traces of (B) B. subtilis wild type and (C) ΔyugO strain treated with IAA-94 chloride channel blocker and upon administration of MTP2 at 5 μg/mL. Membrane potential variation in (D) B. subtilis wild type and (E) ΔyugO strain treated with IAA-94 chloride channel blocker and upon administration of MTP2 at 5 μg/mL, under dark conditions and illumination with error bars representing the standard deviation. The ι1V is calculated from the difference between each condition and the untreated strain (vs B. subtilis wild type (C), vs ΔyugO (D) strain respectively), which is reported in transparency (grey).

Finally, we tested whether chloride channels contribute to the MTP2 response by utilizing the chloride channel blocker indanyloxyacetic acid 94 (IAA-94) at a concentration of 1 μM. Wild type cells treated with IAA-94 alone exhibited a slight depolarization of the membrane resting potential (+20 mV), consistent with reduced Cl^-^ influx (**Figure 4 B**). When MTP2 was co-added with IAA-94, the cells showed a mild hyperpolarization in the dark and a light-induced depolarization that was almost indistinguishable from IAA-94 untreated controls (**Figure 4 B, D**). Thus, blocking Cl^-^ channels did not abolish the MTP2-driven membrane potential dynamics: cells still depolarized upon illumination to a similar extent as in the absence of IAA-94.

In contrast, in the ΔyugO mutant co-treated with IAA-94 and MTP2 (**Figure 4C,E**), we observed a pronounced depolarization in the dark (>40 mV), followed by only a small, persistent hyperpolarization upon illumination (≈ −6 mV). This polarity inversion relative to the canonical response (dark hyperpolarization and light-induced depolarization) is consistent with a major re-weighting of the effective bioelectric circuit: removing a key K^+^ buffering pathway (YugO) while pharmacologically inhibiting IAA-94–sensitive Cl^-^ conductances severely constrains the cell’s ability to counterbalance additional ionic perturbations. In this compromised background, the dominant effect is a shift of the effective leak balance toward a depolarized operating point (depolarized fixed point), reflecting loss of compensatory homeostatic mechanisms rather than a distinct mode of MTP2 action.

Illumination therefore does not invoke an alternative electrophysiological pathway; instead, it partially offsets the depolarizing load present under these conditions, allowing the membrane potential to relax modestly toward a more polarized state. The persistence of this light-induced shift supports a slow homeostatic rebalancing of ionic gradients and/or effective conductances, rather than a response limited by the ultrafast photophysics of MTP2.

Overall, these results support an interconnected regulatory network in which Cl^-^-dependent pathways modulate the MTP2 response and become critical when the primary K^+^-dependent feedback is absent. More broadly, they highlight the potential for bacterial ion homeostasis to reconfigure under combined genetic and pharmacological constraints, underscoring the need for deeper functional characterization of bacterial chloride pathways, many of which remain poorly understood. A summary of all electrophysiological responses in *B. subtili*s WT and ΔyugO, with and without IAA-94, under dark and illuminated conditions is provided in **Table 1**.

**Table 1.** Summary of the electrophysiological (Membrane potential) effects driven by MTP2 (5 μg/mL) in B. subtilis WT and mutant strain, under dark and light (470 nm, 10 s) conditions.

| | Wild type | $\Delta yugO$ | |
| --- | --- | --- | --- |
| **MTP2** | - 86 mV | - 90 mV | **Dark** |
|  | - 75 mV | - 62 mV | **Light** |
|  | K<sup>+</sup> efflux / Cl<sup>-</sup> influx | Reduced K<sup>+</sup> efflux buffering |  |
| **MTP2**<br>**IAA-94** | - 94 mV | - 34 mV | **Dark** |
|  | - 84 mV | - 40 mV | **Light** |
|  | K<sup>+</sup> efflux / reduced Cl<sup>-</sup> conductance | Reduced Cl<sup>-</sup> conductance / reduced K<sup>+</sup> buffering |  |

### 2.6 Role of Active transport

To gain a deeper understanding on how MTP2 perturbs ion transport across the membrane, we challenged the system by partially dissipating the proton motive force (PMF) with the protonophore Carbonyl cyanide 3-chlorophenylhydrazone (CCCP). When supplemented at 10μM, CCCP alone caused a modest depolarization of the resting membrane potential (**Figure 5A**), consistent with increased proton permeability and reduced PMF. Strikingly, however, CCCP did not suppress the MTP2-driven voltage modulation. Instead, co-application of MTP2 in the dark drove the membrane to a markedly hyperpolarized state (≈ −120 mV), exceeding the hyperpolarization observed with MTP2 alone, and a 470 nm light pulse still elicited a robust depolarization with an amplitude nearly twice that measured in untreated cells (**Figure 5A**).These results indicate that MTP2 can modulate bacterial electrophysiology even when PMF-dependent energetics are perturbed, arguing against a mechanism that strictly requires intact ATP synthesis or normal metabolic activity. Rather than simply “removing” voltage, CCCP reshapes the effective homeostatic background, altering the balance of leak currents and compensatory transport and, consequently, the electrical sensitivity to a given MTP2-driven bias. In this pump–leak perspective, CCCP shifts the resting set point while reducing the effective buffering (“clamping”) of *V_m_*, thereby unmasking larger voltage excursions arising from MTP2-mediated perturbations of passive ionic fluxes and/or interfacial ion partitioning. A collective representation of the responses for each experimental condition is shown in **Figure 5B**.

**Figure 5.**
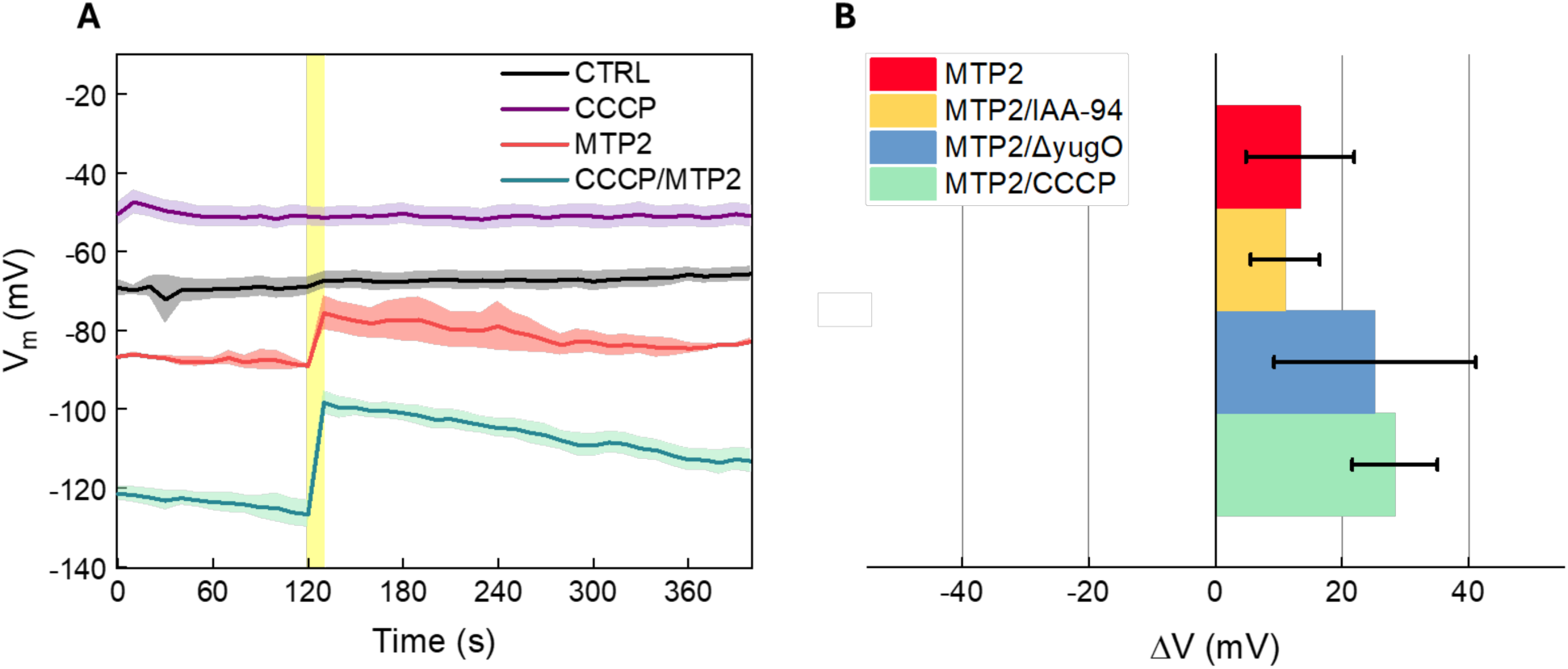
MTP2 effect on K^+^ active transport. (A) Electrophysiological traces of B. subtilis treated with CCCP, hence lacking active transport pathways for homeostasis regulation. (B) Light-induced membrane potential variation for each condition, with error bars representing the standard deviation. For each condition, 9 samples were analysed divided on three different groups from three independent experiments conducted on different days.

### 2.7 Modulation of antibiotic sensitivity

Antibiotic efficacy on bacteria is sensitive to changes in cell membrane potential.^19,46–49^ Thus, here we tested whether our non-genetic optostimulation approach might modulate cell viability at sublethal kanamycin concentrations (found to be 15 µg·mL^-1^ and less).^19^ At 5 µg/mL MTP2 (within the low toxicity range of MTP2 with *B. subtilis* as discussed above in **Figure 1B**), we measured the growth of *B. subtilis* cells at a range of kanamycin concentrations from 0 to 15 µg/mL, with and without illumination at 470 nm in cycles of 10 minutes light, followed by 20 minutes in the dark, repeated throughout the 8h growth. Low kanamycin concentrations 0 to 5 µg/mL showed little impact on cell growth, nor a significant difference due to light or MTP2 (**Figure 6A**, **Figure S5**). At high concentrations of kanamycin 20 µg/mL and above, growth was significantly slowed and became too sporadic to discern any differences due to MTP2 or illumination. However, at intermediate concentrations of kanamycin 10 to 15 µg/mL, we observed an interesting effect of MTP2. The presence of MTP2 in the dark decreased the total cell growth, primarily resulting in a lower final optical density (**Figure 6A**, **Figure S5**. Illumination of *B. subtilis* in the presence of MTP2, meanwhile, restored cell growth to the same levels as seen in the samples without MTP2. **Figure 6B** shows the kanamycin concentration dependence of the MTP2-induced percent variation in area under the growth curve in the dark, compared to light conditions which eliminate this difference. Full data sets showing the areas under the growth curves for each condition are shown in **Figure S6**.

**Figure 6.**
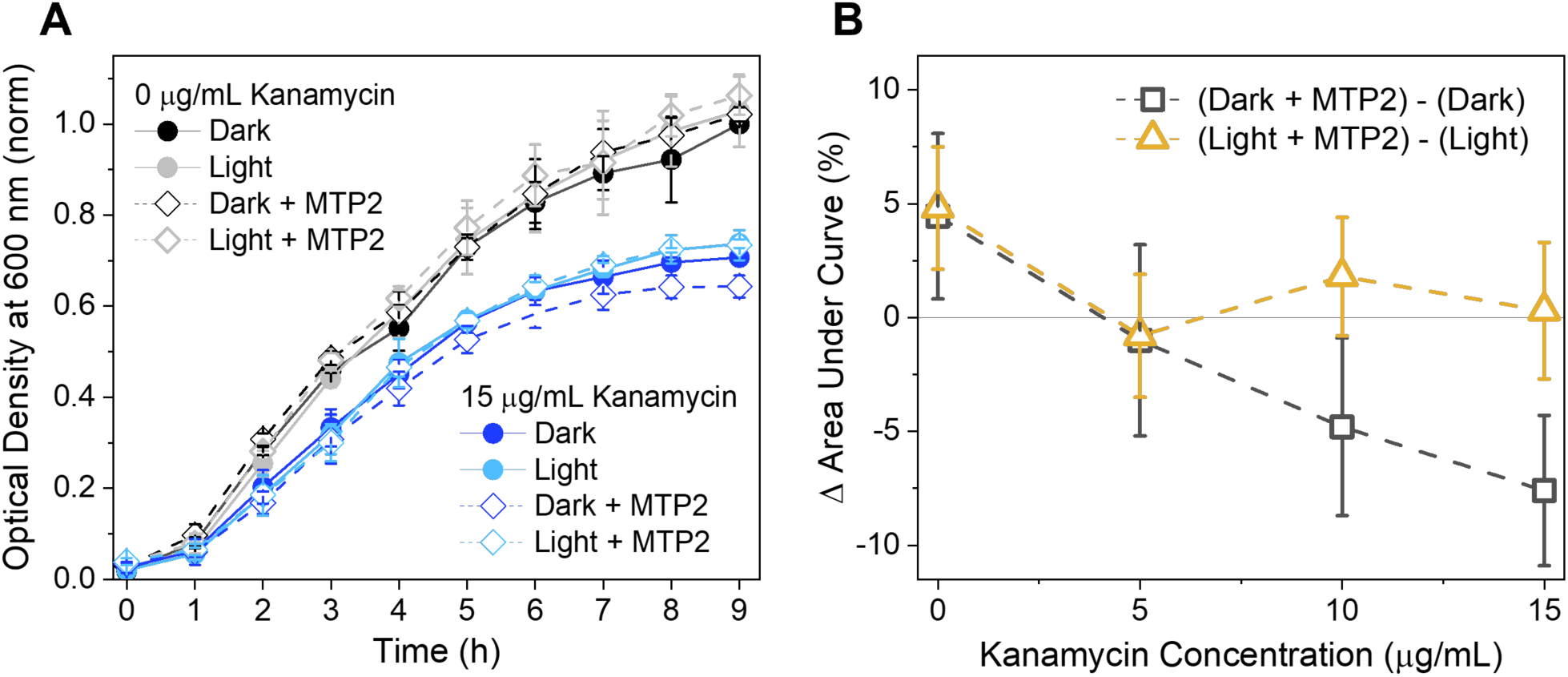
Modulation of antibiotic uptake. (A) Growth of B. subtilis cells as monitored by optical density, showing the impact of MTP2 and 470 nm illumination (light) at 0 and 15 µg/mL kanamycin. (B) The percentage of MTP2-induced change in the area under the B. subtilis growth curves as a function of kanamycin concentration dependence, obtained from subtracting the area under the curves without MTP2 from the curves with MTP2, for both light and dark conditions.

The increase in lethality of kanamycin with *B. subtilis* caused by MTP2 under dark conditions is consistent with prior literature which has established that bacterial uptake of aminoglycoside family of antibiotics, of which kanamycin is part, depends on the polarization state of the cell’s membrane.^17,46–49^ Therefore, in the dark, MTP2-induced hyperpolarization (**Figure 3**) leads to an increased kanamycin uptake at the single cell level, and the observed slowed cell growth of the whole population (**Figure 6A, B**). The restoration of cell growth upon illumination with MTP2, then, implies that the light-induced depolarization due to MTP2 (**Figure 3**) is enough to measurably decrease kanamycin uptake and its bactericidal impact. These results suggest that tailored membrane-targeted, photoswitchable molecules such as MTP2 can modulate antibiotic susceptibility.

## 3. Discussion

Mechanistically, our data support a picture in which MTP2 acts as a phototunable perturbation to the membrane’s effective electrical balance, rather than as a fast, purely photophysical actuator. Although trans–cis isomerization occurs on ultrafast timescales, the electrophysiological response evolves over seconds to minutes, indicating that the measured *V_m_* reflects slow electrical and homeostatic dynamics (ion flux, redistribution, and buffering by endogenous transport) of cells. A useful starting point is charge–capacitance bookkeeping: because the membrane behaves as a capacitor, changes in voltage satisfy Δ*Q* = *C* Δ*V* (or equivalently Δ*V* = Δ*σ*/*C_m_* , where Δ*σ* is the net charge imbalance per membrane area and *C_m_* is the specific capacitance per unit area). Importantly, *C_m_* is not expected to differ strongly between lipid membranes; thus, differences in response amplitude across cell types are more naturally attributed to differences in the effective interfacial perturbation (Δ*σ*) and in how strongly endogenous conductances clamp voltage. In bacteria, relatively small net ionic imbalances can shift *V_m_* by tens of millivolts, and our uptake/ζ-potential measurements indicate that the Gram-positive envelope can electrostatically pre-concentrate cationic amphiphiles, plausibly increasing the effective surface density of MTP2 and thereby its interfacial bias (see SI for quantitative estimates and a comparison with mammalian cells). To formalize this intuition, we describe the voltage dynamics with an effective pump–leak model (detailed in the SI), in which the membrane potential obeys

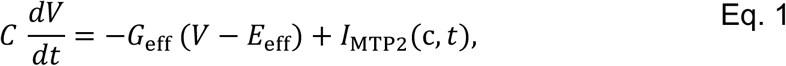

where *G*_eff_ and *E*_eff_ summarize the net background conductances and reversal potentials of the endogenous “bioelectric circuit,” and *I*_MTP2_(*t*) represents the net bias introduced by MTP2. In this framework, MTP2 need not be assigned to a single ion species; instead, it shifts the balance of currents in a photo-state- and context-dependent manner. Circuit principles then immediately provide scaling relationships: the steady-state shift satisfies Δ*V* ∼ *I*_MTP2_/*G*_eff_, while the fast electrical timescale scales as *τ* ∼ *C*/*G*_eff_ . The long-lived plateaus and slow drifts observed experimentally are naturally captured by allowing the effective bias and/or effective leak balance to evolve on slow timescales reflecting ion homeostasis and redistribution, rather than the ultrafast MTP2 photophysics.

Within this unified electrical description, the wild type response can be interpreted as follows. In the dark, MTP2 shifts the steady state to a more negative *V_m_* (hyperpolarization), consistent with a bias that favours net outward positive charge movement and/or suppresses depolarizing leak components. Upon illumination, the MTP2 photo-state shifts the current balance in the opposite direction, producing depolarization. The persistence of the post-illumination relaxation over hundreds of seconds indicates that recovery is governed by slow re-equilibration of ionic gradients and active transport, rather than by cis-to-trans back-relaxation in solution. In other words, MTP2 acts as an optical trigger that repositions the effective electrical set point, while endogenous transport processes determine the observed kinetics and waveform.

The mutant experiments further demonstrate that the MTP2 response is strongly shaped by which native conductances are available to buffer voltage perturbations. In ΔyugO, the dark hyperpolarization remains comparable to wild type, indicating that the resting shift induced by MTP2 does not require YugO specifically and can be supported by other K^+^ routes and/or non-specific background conductances. By contrast, the light-induced depolarization is strongly amplified in *ΔyugO*, consistent with YugO functioning primarily as a negative-feedback K^+^ shunt that clamps departures from the resting potential. In the pump–leak picture, deleting YugO effectively reduces the stabilizing conductance available during depolarizing excursions (decreasing *G*_eff_ in the relevant operating regime), thereby increasing Δ*V* for the same MTP2-driven bias.

The behaviour under combined genetic and pharmacological constraints underscores that the bacterial voltage set point can be qualitatively reconfigured. In wild type cells, IAA-94 modestly depolarizes the resting potential, but co-application of MTP2 preserves the characteristic light-evoked depolarization, indicating that IAA-94–sensitive chloride pathways are not strictly required for the canonical MTP2 response. In striking contrast, in the ΔyugO background co-treated with IAA-94 and MTP2, cells adopt a strongly depolarized dark steady state and show a small, persistent hyperpolarization upon illumination. In the circuit framework, this sign inversion indicates that removing both a dominant K^+^ buffering pathway (YugO) and IAA-94–sensitive Cl^-^ conductance places the system in a different operating regime where the effective background balance *E*_eff_ and/or *G*_eff_ is substantially altered, and the MTP2 photo-switch now partially relieves (rather than drives) the dominant bias. The persistence of the light-induced hyperpolarizing shift further supports the interpretation that the light pulse switches the system into a long-lived electrophysiological state, consistent with slow re-equilibration of gradients and transport in a constrained homeostatic network.

Consistent with this view, CCCP experiments show that perturbing energetics does not abolish the MTP2 response but can instead amplify it. CCCP alone depolarizes *V_m_*, as expected from increased proton permeability and partial dissipation of PMF. However, CCCP plus MTP2 yields a markedly deeper dark hyperpolarization and a larger light-evoked depolarization. In the pump–leak formulation, this behaviour is naturally explained if CCCP reshapes the effective background circuit, shifting the resting set point and modifying the degree of voltage “clamping”, so that a given MTP2-driven bias produces a larger Δ*V*. Thus, CCCP does not simply remove voltage; it alters the homeostatic landscape in which MTP2 operates, revealing how strongly the observed response depends on the availability of compensatory transport pathways.

Finally, the same electrical framework provides a consistent interpretation of the antibiotic proof-of-concept experiments. Aminoglycoside uptake is sensitive to membrane polarization; therefore, the MTP2-driven hyperpolarized states are expected to favour uptake, while light-driven depolarization should attenuate it. In this sense, MTP2 enables external control over an electrophysiologically coupled phenotype, linking optical perturbations of the effective membrane “circuit” to downstream functional outcomes.

Overall, our results emphasize that bacterial membrane voltage is an emergent property of a tightly coupled network of passive permeabilities and active transport processes, and that optically controlled membrane-targeted amphiphiles such as MTP2 can probe and reconfigure this network. The ability of *B. subtilis* to buffer, amplify, or even invert the sign of the response depending on genetic and pharmacological context highlights a high degree of electrophysiological plasticity, consistent with the view that microbes dynamically reallocate ionic pathways to preserve homeostasis under perturbation.

These findings might have far-reaching applications. In bacteria, light-driven modulators could non-invasively tune biofilm formation, electrical signaling, or antibiotic uptake. More broadly, the same chemical photoswitch strategy could be applied to probe bioelectric phenomena in diverse biological contexts: from wound healing,^50^ to plant physiology,^51^ cells extrusion^52^ and cancer development. In this latter case, bioelectricity has been seen to play an important role in cell proliferation and metastasis.^53,54^ By providing a non-genetic, reversible handle on voltage, MTP2 and related dipolar switches may enable new approaches for fundamental bioelectric research.

## 4. Materials and methods

### 4.1 Growth conditions, viability assay and preparation of agarose pads for Bacillus subtilis

Strains used in this work are the following: NCIB 3610 wild type and NCIB 3610 *yugO::neo*. The following procedure for culturing and stock preparation has been employed for all the strains forementioned. Glycerol stock was set up using 50% of culture broth and, to avoid crystal formations upon freezing, 50% glycerol. Strains were then stored at -80°C. Cells were then streaked on lysogeny-broth (LB) 1.5% agar and incubated overnight in a 37°C non-shaking incubator. A single colony was picked from this plate, inoculated in liquid LB and incubated at 37°C shaking (200 rpm) overnight.

MTP2 was prepared according to a previously reported synthetic protocol: all reaction conditions, purification steps, and characterization methods were performed as outlined by Sesti et al.^35^ To assess the toxicity of MTP2, and for the antibiotic modulation experiments with kanamycin, cells were cultured as described above and then diluted up to OD_600_ = 0.1. For the viability assay, MTP2 was diluted in Milli-Q sterile water and added to the diluted cell suspensions using increasing concentrations, respectively 0 (control), 5, 10, 20 and 50 μg/mL. For the antibiotic modulation experiments, 0 or 5 μg/mL MTP2 and 0, 5, 10, 15, or 20 μg/mL kanamycin were added to the cell suspensions. Cells were incubated for 10 minutes with the molecules in a 24 multi-well plate containing 1 or 2 mL of culture in each well. Subsequently, this plate was placed on the OptoBiolabs optoWELL^®^ which allows precise light stimulation with uniform spatial distribution of light and high temporal resolution. Light stimulation was carried out for 8 to 9 hours with a pattern consisting of 10 minutes of 470 nm light exposure and 20 minutes of dark cycle. The whole assay was carried out at 37°C in a shaking incubator (160 rpm). To monitor cellular growth, optical density was measured, at the wavelength of 600 nm, every hour using a Tecan Spark 10M plate reader. For microscopy assay, overnight culture in liquid LB were pelleted and washed once with resuspension media (RM; composition per 1 litre: 46 μg FeCl_2_, 4.8 g MgSO_4_, 12.6 mg MnCl_2_, 535 mg NH_4_Cl, 106 mg Na_2_SO_4_, 68 mg KH_2_PO_4_, 96.5 mg NH_4_NO_3_, 219 mg CaCl_2_, 2 g monosodium L-glutamate), and then incubated in RM at 37°C shaking for an hour prior to microscopy assay. Following incubation with RM, cells were deposited on RM 1.5% weight/volume Low Melting Point (LMP) agarose pads prepared as described previously.^25^ When specified, the Nernstian dye and Push-Pull were added at the following concentrations: TMRM at 100 nM (Molecular Probes); MTP2 at 2, 5 and 10 µg/mL, as described in the Results section, IAA94 at a concentration of 1μM and CCCP at a concentration of 10μM.

### 4.2 MTP2 cellular uptake measurement

Cells suspended in LB medium (OD_600_ = 1) were incubated with 10 µg/mL MTP2 at 25°C for 1 hour. The illuminated samples (470 nm) were treated using the following illumination protocol using a 470nm Thorlabs LED (M470L5): 10 min of illumination followed by 20 min in dark, repeated. The samples were then centrifuged, and the supernatant was transferred to 1-cm cuvettes for absorbance measurements with a SHIMADZU UV-1900i UV-Vis spectrophotometer. Control samples without bacteria were treated with this same protocol and were used to obtain the pre-uptake absorbance of MTP2.

### 4.3 Transient absorption spectroscopy

The measurements were carried out in transmission mode using a regenerative amplified Ti:Sapphire laser system from Coherent (operating at 1 kHz, with a fundamental wavelength of 800 nm). The laser beam is split into two parts: the pump, a narrowband beam (∼10 nm) generated via optical parametric amplification using a beta barium borate (BBO) crystal, and the probe, a broadband white-light continuum (400-750 nm) produced with a CaF2 crystal. The pump beam excites the sample at 520 nm, while the probe beam monitors the induced changes in the excited region. After passing through the sample, the probe is directed into a monochromator and detected by a CCD camera. The signal measured is ΔT/T where ΔT = T_on_ – T_off_; T_on_ and T_off_ represent the transmitted probe intensities with the pump beam turned on and off, respectively, modulated by an optical chopper operating at 0.5 kHz. The TAS maps are acquired by adjusting the temporal delay between the pump and probe beams using a motorized delay stage, allowing for delays up to 1.25 ns.

### 4.4 Time-lapse microscopy and light stimulation

For time-lapse and 470 nm light stimulation experiments, the fluorescence microscope Leica DMi8, equipped with a Leica K5 camera and an objective lens HC PL APO 100x/1.40 OIL PH3, was used. TMRM fluorescence was detected with 150 ms exposure with a custom filter cube FITC/TRITC, size P, Ex 470/40, 548/15, dichroic 495, 562 and Em 515/30, 590/45 (Leica). For 470 nm stimulation the same cube was used with 10 seconds exposure. The light power of the 470 nm stimulation was measured with the PM400 power meter (Thorlabs) equipped with the S175C sensor (Thorlabs) and the power density calculated in accordance with the area of the field of view.

For each condition, 9 samples were analysed divided on three different groups from three independent experiments conducted on different days. Time-lapse duration was 2 minutes before 470 nm stimulation, with acquisition interval of 10 seconds. Immediately after, another 5 minutes time-lapse with same acquisition interval was conducted, where 470 nm exposure occurred once after the first TMRM image acquisition. For the 30-min time-lapse instead the first interval was 2 minutes before light stimulation and acquisition interval every 10 seconds but the second time-lapse lasted for 30 minutes with a light pulse of 10 seconds every 10 minutes.

### 4.5 Membrane potential estimation

Estimation of B. subtilis membrane potential in millivolt was performed as described by Ehrenberg et al.,^55^ using the following equation:

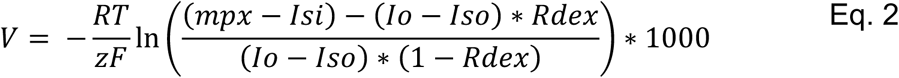

Where R is the gas constant, T is the temperature in Kelvin, z is the charge of the dye, F is the Faraday constant, mpx is the integrated density from analysed cells, Io is the background intensity, Isi is the autofluorescence of the cell (measured from cells not exposed to TMRM), Iso is the background intensity in the absence of TMRM. Calculations were performed with Python.

### 4.6 **ζ** potential measurement

Samples were diluted up to OD_600_=0.5, centrifuged and resuspended in deionized water and MTP2 at concentrations of 5, 10, 30, 50 and 80 μg/ml. Measurements were performed on a Malvern Zetasizer Nano ZS (Malvern Instruments, Malvern, U.K.) at RT. Data points given are an average of 3 biological replicates with 3 measurements each.

### 4.7 OD variation growth assay

Bacteria culture was grown in liquid LB over night at 37° while shaking at 160 rpm. Cells in stationary phase were then diluted up to an optical density at 600 nm = 0.1 and placed into spectrophotometer 1-mL cuvettes and sealed with parafilm. Cells were then incubated at 37° for 7 hr and exposed to a photostimulation cycle which consists in 10 min of light exposure using a 470nm Thorlabs LED (M470L5) and 20 minutes of dark. After 1h and every hour, optical density was measured using a SHIMADZU UV-1900i UV-Vis spectrophotometer at a wavelength of 600 nm.

## 5. Acknowledgements

This work is co-funded by the European Union (ERC, EOS, 101115925). Views and opinions expressed are, however, those of the author(s) only and do not necessarily reflect those of the European Union or the European Research Council. Neither the European Union nor the granting authority can be held responsible for them. MA acknowledges the funding support from the University of Warwick Research Development Fund Strategic Award and the Leverhulme Trust Research Grant (RPG-2024-327).

