## Supplementary Information for "A Dipolar Photoswitch Modulates Bacterial Membrane Potential and Reveals Context-dependent Bioelectrical Circuitry"

### ***Supplementary Information for "A Dipolar Photoswitch Modulates Bacterial Membrane Potential and Reveals Ion-channel Buffering Networks"***

Pietro Bertolotti[1,2], Fabio Marangi[1], Edoardo Cianflone[1,3], Andrea Pianetti[1], Helena R. Keller[1,3], Arianna Magni[4], Valentino Romano[3], Marta Ghidoli[5], Tailise Carolina de Souza-Guerreiro[6], Chiara Bertarelli[5], Guglielmo Lanzani[1,3], Munehiro Asally[6], Giuseppe Maria Paternò[1,3*]

[1] Center for Nanoscience and Technology, Istituto Italiano di Tecnologia, Via Rubattino 81, 20134 Milan, Italy
[2] Department of Electronics, Information and Bioengineering, Politecnico di Milano, Via Ponzio 34/5, 20133 Milan, Italy
[3] Department of Physics, Politecnico di Milano, Piazza Leonardo Da Vinci 32, 20133 Milan, Italy
[4] Department of Materials Science and Engineering, Stanford University, Stanford, CA 94305, USA
[5] Department of Chemistry, Materials and Chemical Engineering, "Giulio Natta" Politecnico di Milano, Piazza Leonardo Da Vinci 32, 20133 Milan, Italy
[6] School of Life Sciences, University of Warwick, Coventry, CV4 7AL, UK

#### S1. Estimation of MTP2 molecules per B. subtilis cell (cell-associated upper bound).

We estimated the number of MTP2 molecules associated with a single B. subtilis cell from the depletion/retention assay performed at OD600 = 1, using a mass-balance calculation. This estimate should be interpreted as an upper bound on membrane-inserted molecules because the pelleted fraction can include envelope-/wall-bound pools in addition to molecules partitioned into the cytoplasmic membrane.

**Inputs:**

- Initial concentration: $c_0 = 10\ \mu g\cdot mL^{-1}$
- Retained fraction: $f = 0.45$
- Molar mass: $MW = 714\ g\cdot mol^{-1}$
- Cell density at OD600 = 1: $N_{cells} \approx (1\text{–}10)\times 10^8\ cells\cdot mL^{-1}$

**Calculation (per 1 mL of suspension):**

$m\_ret = f \cdot c_0 = 0.45 \times 10\ \mu g\cdot mL^{-1} = 4.5\ \mu g\cdot mL^{-1}$

$n\_ret = m\_ret / MW = (4.5\times 10^{-6}\ g) / (714\ g\cdot mol^{-1}) = 6.30\times 10^{-9}\ mol$

$N\_ret = n\_ret \cdot N\_A = 6.30\times 10^{-9}\ mol \times 6.022\times 10^{23}\ mol^{-1} = 3.80\times 10^{15}\ molecules\cdot mL^{-1}$

Dividing by the number of cells per mL gives the number of cell-associated molecules per cell:

$N\_cell = N\_ret / N\_cells$

**Representative values:**

| Assumed Ncells (cells·mL$^{-1}$) | Estimated Ncell (molecules·cell$^{-1}$) |
|---|---|
| 1e+08 | 3.80e+07 |
| 3e+08 | 1.27e+07 |
| 5e+08 | 7.59e+06 |
| 1e+09 | 3.80e+06 |

Across plausible OD-to-cell conversions, this corresponds to ~$4\times10^{6}$ to ~$4\times10^{7}$ MTP2 molecules per cell at 10 μg·mL$^{-1}$ and 45% retention.

<u>Order-of-magnitude surface density (upper bound):</u> Approximating B. subtilis as a cylinder of radius r = 0.5 μm and cylinder length L = 2.5 μm with two hemispherical caps, the surface area is $A \approx 2\pi rL + 4\pi r^2$ = 11.0 μm$^2$ (~1.1e+07 nm$^2$). The corresponding upper-bound areal density at 10 μg·mL$^{-1}$ is ~0.3–3 molecules·nm$^{-2}$, indicating that a substantial fraction of the retained MTP2 likely resides in envelope-associated pools rather than as a uniformly distributed in the cytoplasmic membrane.

**S2. Charge–capacitance bookkeeping: ions required for a given ΔVm**

The membrane behaves electrically as a capacitor. For a voltage change ΔV, the required net charge imbalance satisfies:

$\Delta Q = C \cdot \Delta V$ and $\Delta V = \Delta\sigma / C\_m$

where C is the total membrane capacitance, C_m is the specific capacitance (per unit area), and Δσ is the net charge imbalance per unit area. For lipid membranes, C_m is typically ~1 μF·cm$^{-2}$ (≈ 0.01 F·m$^{-2}$)[1]; thus, differences in response amplitude across cell types are generally driven by differences in Δσ and/or effective conductance 'clamping', rather than large differences in C_m.

---

[1] Treat the membrane as a parallel-plate capacitor:

$$C_m = \frac{\varepsilon_r \varepsilon_0}{d}$$

- $\varepsilon_0 \approx 8.85 \times 10^{-12}$ F/m
- relative permittivity of the hydrophobic core $\varepsilon_r \approx 2 - 4$
- membrane thickness $d \approx 4 - 5$ nm $= 4 - 5 \times 10^{-9}$ m

Plugging in typical values gives $C_m$of order $\sim 0.01$ pF/$\mu$m$^2$. This is confirmed by experiments in HEK and estimation in bacteria (Annu Rev Biophys. 2024 Jul;53(1):487-510.)

**Geometric and electrical parameters used for order-of-magnitude estimates:**

| Cell type | Surface area A | Total capacitance C (=C_m·A) | Volume V |
|---|---|---|---|
| B. subtilis (cylinder + caps; r=0.5 μm, L=2.5 μm) | 11.0 μm² | 1.10e-13 F | 2.49 fL |
| HEK293 (sphere; r=10 μm) | 1257 μm² | 1.26e-11 F | 4.19 pL |

**Ions required for a 20 mV voltage shift:**

B. subtilis: C = 1.10e-13 F → ΔQ = C·ΔV = 2.20e-15 C → N ≈ 1.37e+04 monovalent ions

HEK293: C = 1.26e-11 F → ΔQ = C·ΔV = 2.51e-13 C → N ≈ 1.57e+06 monovalent ions

For B. subtilis, this corresponds to a concentration change of ~9.2 μM in a ~2.49 fL volume; for HEK293, the analogous change is ~0.62 μM in a ~4.19 pL volume.

Voltage change in HEK293 produced by the same net ion number that yields 20 mV in B. subtilis:

Using N = 1.37e+04 ions → ΔV_HEK = (N·e)/C_HEK = 0.17 mV

Thus, an identical net ion number that produces a clear ~20 mV shift in B. subtilis would yield only ~0.1–0.2 mV in a HEK293-sized cell, i.e., at or below the noise level of most optical voltage measurements. This comparison is intended as order-of-magnitude estimation rather than a claim about identical molecular loading across systems.

**S.3. Minimal pump–leak model of MTP2-induced voltage changes.** To rationalize the order of magnitude of the MTP2-induced voltage responses, we used a single-compartment pump–leak model for a B. subtilis cell. The membrane potential $V$obeys

$$C\,\frac{dV}{dt} = -G_{\text{leak}}\,(V - E_{\text{leak}}) + I_{\text{MTP2}}(t,c) \quad \text{eq. S1}$$

Here, $G_{\text{leak}}$ and $E_{\text{leak}}$ are effective parameters that lump together the background permeabilities and homeostatic transport operating around the resting state; the goal is not to attribute the response to a single ion species, but to capture how a perturbation is converted into a voltage waveform by the endogenous electrical “circuit”. We implement $I_{\text{MTP2}}(t,c)$ as a two-state (dark vs illuminated) current whose amplitude depends on MTP2 concentration $c$. During the 10 s blue-light pulse, the current switches from a dark value $I_{\mathrm{dark}}(c)$ to an illuminated value $I_{\mathrm{light}}(c)$. After the pulse, the bias current relaxes back toward $I_{\mathrm{dark}}(c)$ with a slow recovery time constant $\tau_{\mathrm{rec}}$(hundreds of seconds) to phenomenologically represent slow re-equilibration of ionic gradients and transport activities. Because TA measurements show that MTP2 photochemistry is ultrafast (ps photoisomerization; $\sim 10^2$ μs back-relaxation), $I_{\mathrm{MTP2}}$ can be treated as switching effectively instantaneously on the seconds-to-minutes timescale of the electrophysiological recordings.

In this model, the steady-state voltage shift scales as $\Delta V \approx I_{\mathrm{MTP2}}/G_{\mathrm{leak}}$, while the fast electrical time constant is $\tau = C/G_{\mathrm{leak}}$. Slow post-stimulus plateaus and drifts arise from the slow evolution of $I_{\mathrm{MTP2}}$ (or, equivalently, slow changes in the effective leak balance) rather than from MTP2 photophysics.

For simplicity, we assume a linear concentration–effect relation for the bias current over 0–10 μg$\cdot$ $\mathrm{mL}^{-1}$, $I_{\mathrm{dark/light}}(c) \propto c$. Under this approximation, the model predicts a small dark shift at 2 μg$\cdot$ $\mathrm{mL}^{-1}$ that is not resolved experimentally; we attribute this discrepancy to a combination of (i) non-linear dose–response in effective interfacial loading and channel recruitment, and (ii) finite sensitivity/noise in the optical voltage readout at low amplitudes, rather than to a failure of the pump–leak framework itself. Importantly, this deviation at 2 μg$\cdot$ $\mathrm{mL}^{-1}$ does not affect the main conclusions drawn from the model, which are based on the order of magnitude of the MTP2-induced ion fluxes and the tens-of-millivolts responses observed at higher concentrations. The linear concentration dependence should therefore be viewed as a convenient phenomenological approximation that interpolates between the experimentally constrained endpoints (no effect at 0 μg$\cdot$ $\mathrm{mL}^{-1}$ and strong hyperpolarization/depolarization at 10 μg$\cdot$ $\mathrm{mL}^{-1}$), rather than as a precise description of the microscopic dose–response relation.

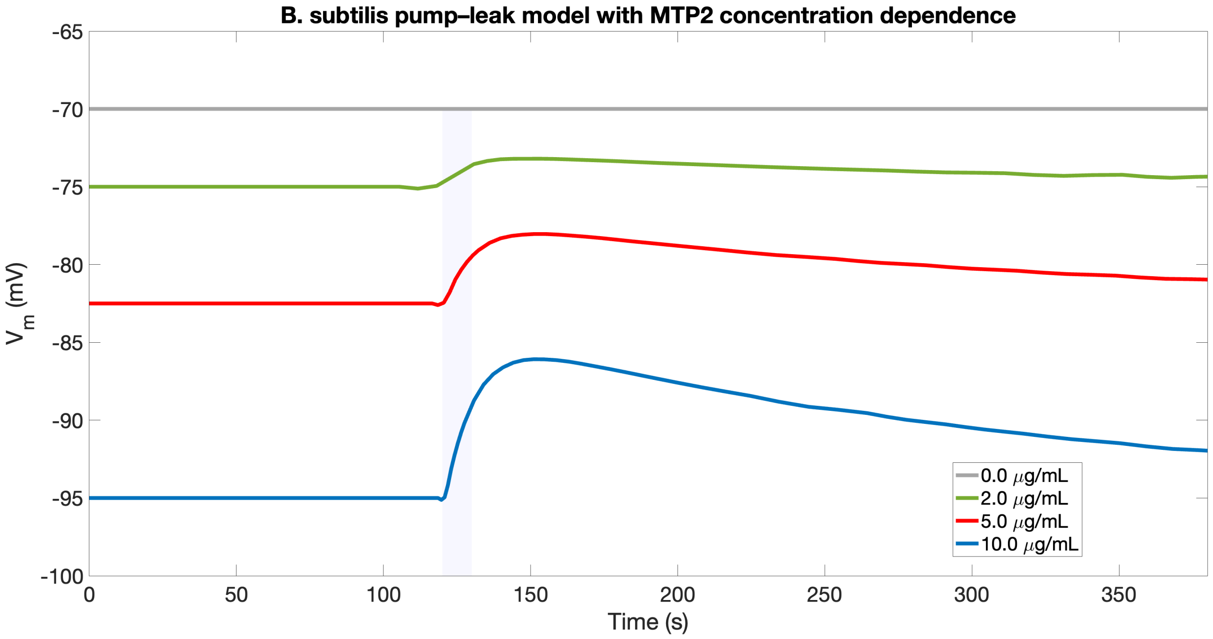


***Figure S1:*** *Simulated voltage changes upon increase of MTP2 concentration.*

**Pump–leak modelling of the ΔyugO strain.** To explore how the voltage-gated $K^+$ channel YugO shapes the MTP2 response, we extended the single-compartment pump–leak model used for wild-type *B. subtilis* to describe three conditions: WT + MTP2 (5 μg·mL⁻¹), ΔyugO control (no MTP2), and ΔyugO + MTP2 (5 μg·mL⁻¹). In all cases, the membrane potential $V$ obeys the same pump-leak equation. The baseline leak conductance was again chosen such that the fast electrical time constant $\tau = C/G_{\text{leak}}$is ≈ 10 s, and a slower recovery of the MTP2 bias with time constant $t_{\text{MTP2}} \sim 200\text{s}$ was included to reproduce the long relaxation observed after the light pulse (see previous section).

For each genotype and condition, the MTP2 term was calibrated via steady-state constraints. For WT + MTP2 at 5 μg·mL⁻¹ we imposed a dark steady state $V_{\text{dark}}^{\text{WT}} \approx -90$ mV and a light plateau $V_{\text{light}}^{\text{WT}} \approx -79$ mV (ΔV ≈ 11 mV), using the relation $V_{\text{ss}} = E_{\text{leak}} + I_{\text{MTP2}}/G_{\text{leak}}$ to determine $I_{\text{dark}}$ and $I_{\text{light}}$ from a common $E_{\text{leak}}^{\text{WT}}$. The ΔyugO control was represented by the same $C$ and $G_{\text{leak}}$, but with a slightly depolarized leak reversal $E_{\text{leak}}^{\Delta\text{yugO}} \approx -80\text{mV}$ and $I_{\text{MTP2}} \equiv 0$, yielding a stable resting potential around −80 mV with no light response. For ΔyugO + MTP2 at 5 μg·mL⁻¹ we kept $E_{\text{leak}}^{\Delta\text{yugO}}$ fixed and chose the MTP2 bias such that the dark steady state shifted to $V_{\text{dark}}^{\Delta\text{yugO}} \approx -90$ mV (hyperpolarized relative to the ΔyugO control) and the light plateau reached $V_{\text{light}}^{\Delta\text{yugO}} \approx -60$ mV (ΔV ≈ 30 mV), matching the experimentally observed larger depolarization amplitude in the mutant.

Within this unified framework, the key difference between WT and ΔyugO is not the presence or absence of dark hyperpolarization (both strains hyperpolarize in the presence of MTP2), but the effective conductance that opposes depolarization during light. In a steady-state approximation, the light-induced voltage change is $\Delta V \approx I_{\mathrm{MTP2}}/G_{\mathrm{eff}}$ . In wild type, depolarization opens YugO and increases the effective K$^{+}$ conductance, such that $G_{\mathrm{eff}}^{\mathrm{WT}} = G_{\mathrm{bg}} + G_{\mathrm{YugO}}$ and the same MTP2-driven current produces a relatively lower depolarization (≈11 mV). In the ΔyugO strain, the YugO-dependent hyperpolarizing pathway is absent, $G_{\mathrm{eff}}^{\Delta\mathrm{yugO}} \approx G_{\mathrm{bg}}$ is smaller, and therefore the same order-of-magnitude bias yields a much larger voltage excursion (≈ 30 mV). The simulations thus support a picture in which MTP2 can hyperpolarize both WT and ΔyugO cells via its effect on surface electrostatics and background K$^{+}$/Cl$^{-}$ conductances, while YugO primarily acts as a dynamic K$^{+}$ shunt that limits the amplitude of light-evoked depolarization in wild-type cells.

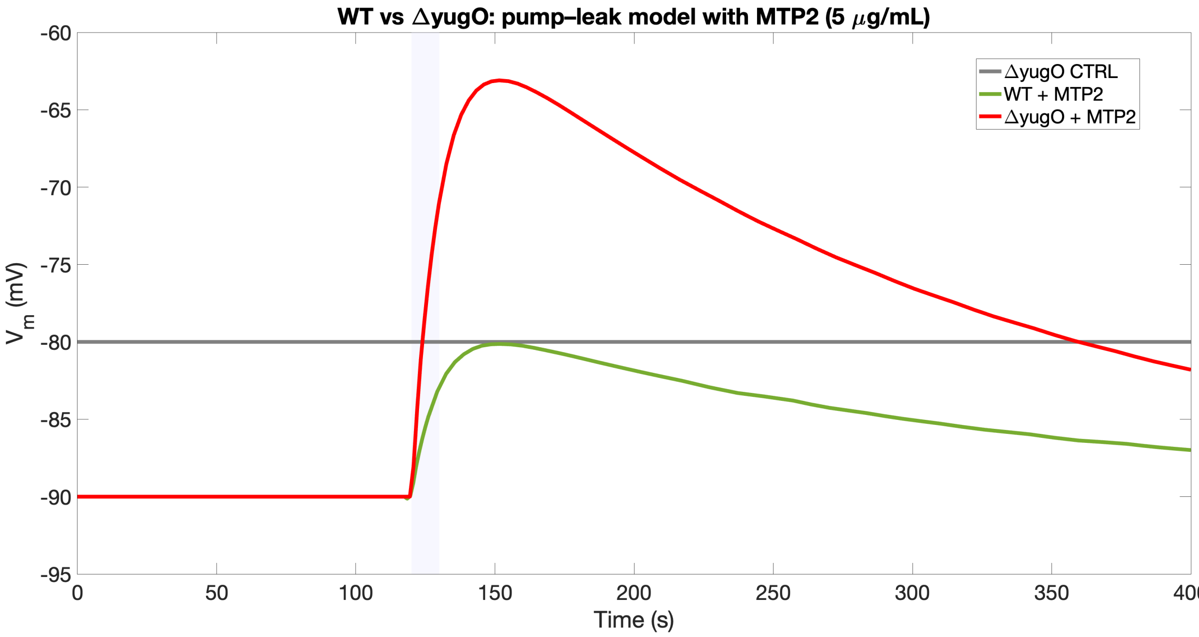


***Figure S2.*** *Simulated voltage changes in the YuGo deleted B. Subtilis strain upon administration of MTP2 at 5 mg/mL. A comparison with the wild-type is also shown.*

**Pump–leak simulations for ΔyugO and pharmacological inhibition of Cl$^{-}$ pathways.** Genetic deletion of YugO and pharmacological inhibition of Cl$^{-}$ pathways were incorporated phenomenologically through changes in $E_{\mathrm{leak}}$ and/or $G_{\mathrm{leak}}$, reflecting altered availability of stabilizing K$^{+}$ and Cl$^{-}$ currents that normally buffer perturbations of $V_m$ . The MTP2 contribution was represented as a two-level input current ($I_{\mathrm{dark}}$ in the dark and $I_{\mathrm{light}}$during illumination; 10 s pulse) whose magnitudes were calibrated directly from the experimental steady-state plateaus via $V_{\mathrm{ss}} = E_{\mathrm{leak}} + I_{\mathrm{MTP2}}/G_{\mathrm{leak}}$. For conditions showing a slow post-pulse drift, $I_{\mathrm{MTP2}}(t)$ was allowed to relax from $I_{\mathrm{light}}$ back towards $I_{\mathrm{dark}}$ with a phenomenological recovery time $t_{\mathrm{recovery}}$ (hundreds of seconds). Conversely, for the ΔyugO + IAA-94 + MTP2 condition, where the voltage remains on a stable plateau with minimal recovery over the full recording window, the model was run in a “persistent state” mode in which the light pulse switches the effective MTP2 bias to a long-lived value (equivalently, $t_{\mathrm{recovery}}$ was taken much longer than the experiment). Within this formalism, the key qualitative findings are naturally reproduced: (i) ΔyugO cells remain close to a baseline set by $E_{\mathrm{leak}}$ in the absence of MTP2; (ii) addition of MTP2 produces a dark shift and a light-evoked change whose amplitude depends on the effective clamping conductance $G_{\mathrm{leak}}$; and (iii) combining ΔyugO with IAA-94 places the system into a markedly different operating regime, in which the available ionic buffering is reduced and the MTP2 bias drives a strongly depolarized dark steady state and an inverted, weak light response. While this model is not intended as a detailed mechanistic reconstruction of specific transporters, it provides a compact description of how removing dominant $K^+$ (YugO) and $Cl^-$ pathways can qualitatively reshape the electrophysiological response to the same molecular perturbation.

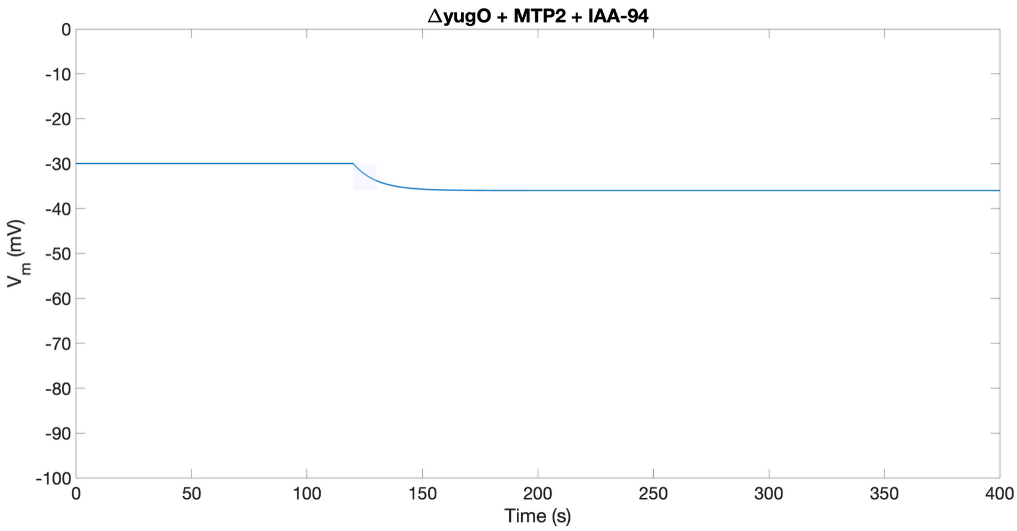


***Figure S3.*** *Simulated voltage changes in the YuGo deleted B. Subtilis strain upon the combinated administration of MTP2 at 5 mg/mL and the pharmacological chloride ion blocker IAA94.*

**Pump–leak simulations incorporating CCCP**. To assess whether the CCCP experiments can be accounted for within a minimal electrical framework, we extended the pump–leak model used throughout this work to explicitly represent CCCP as a perturbation of the effective background electrophysiology.

CCCP was implemented phenomenologically by shifting the effective leak reversal potential to a more depolarized value ($E_{\text{leak}}^{\text{CCCP}} \approx -50$ mV) compared to control ($E_{\text{leak}}^{\text{CTRL}} \approx -65$ to $-70$ mV), reproducing the depolarized baseline observed experimentally in CCCP-treated cells. In addition, to capture the larger voltage excursion observed for CCCP + MTP2 compared with MTP2 alone, we allowed for a reduction of the effective leak conductance under CCCP by a multiplicative factor $\beta_{\text{CCCP}} < 1$ (i.e. $G_{\text{leak}}^{\text{CCCP}} = \beta_{\text{CCCP}} \times G_{\text{leak}}^{\text{CTRL}}$), corresponding to reduced "clamping" of $V_m$ by homeostatic conductances in the CCCP-perturbed state.

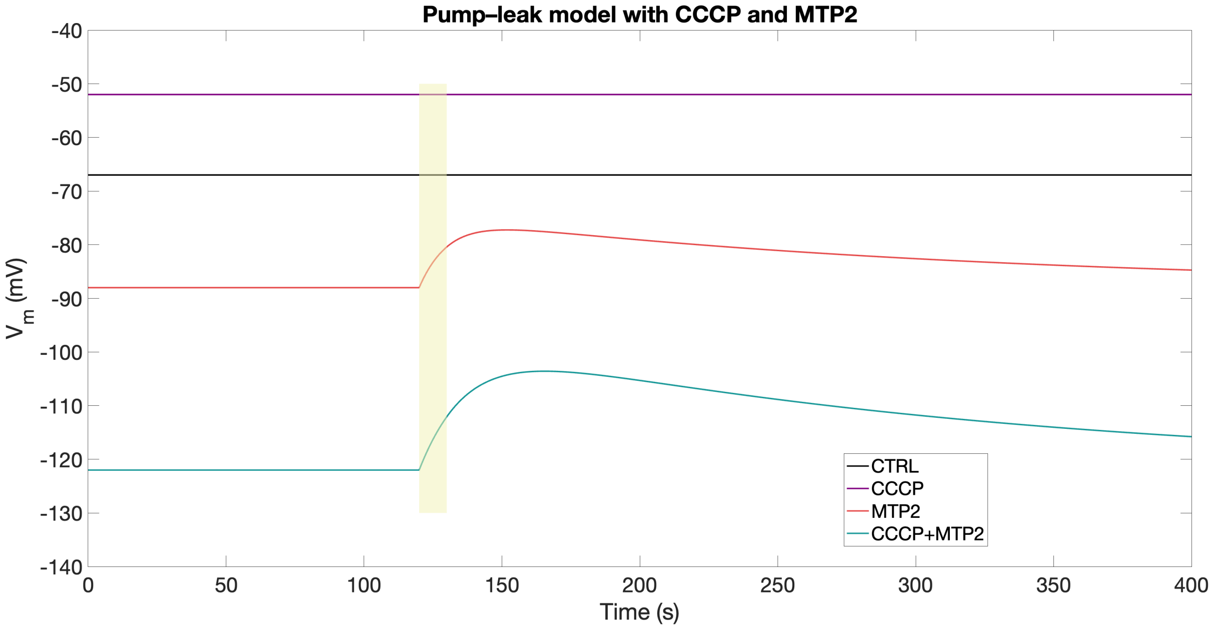


***Figure S4.*** *Simulated voltage changes in the wild-type B. Subtilis strain upon the combinated administration of MTP2 at 5 mg/mL and CCCP.*

The MTP2 contribution was represented as a bias current with two values, $I_{\text{dark}}$ and $I_{\text{light}}$, corresponding to dark and illuminated conditions (10 s pulse), calibrated from the experimentally observed steady-state plateaus through $V_{\text{ss}} = E_{\text{leak}} + I_{\text{MTP2}}/G_{\text{leak}}$. To mimic the slow drift after illumination, $I_{\text{MTP2}}(t)$ was allowed to relax exponentially from $I_{\text{light}}$ back toward $I_{\text{dark}}$ with a phenomenological recovery time constant $t_{\text{recovery}}$ (typically 100–300 s), reflecting slow physiological recovery processes (e.g., ion homeostasis and PMF re-equilibration) that occur on timescales far longer than the ultrafast photophysics of MTP2. With this parameterization, the model reproduces (i) the CCCP-induced depolarization in the absence of MTP2, (ii) dark hyperpolarization and light-evoked depolarization in the presence of MTP2, and (iii) the enhanced hyperpolarization amplitude and larger voltage excursion observed for CCCP + MTP2, consistent with CCCP primarily reshaping the effective leak background and thereby modulating the electrical sensitivity to a given MTP2-driven bias current.

#### S4. MATLAB code

Numerical integration was performed using MATLAB (ode45/ode15s) using Eq. (S1). A representative implementation is reported below. Mutant and pharmacological conditions were simulated within the same formalism by adjusting the effective parameters $G_{\text{leak}}$, $E_{\text{leak}}$, the bias amplitudes $I_{\text{dark}}(c)$, $I_{\text{light}}(c)$, and the recovery constant $\tau_{\text{rec}}$ (Table S1), reflecting changes in the effective background clamp and in the net MTP2-induced current bias.

***Table S1***

| Condition | Parameters changed (relative to WT) | Rationale |
|---|---|---|
| ΔyugO | $G_{\text{leak}}$ ↑and/or $I_{\text{light}}$ ↑ | reduced fast $K^+$ buffering increases net depolarization |
| IAA-94 | $E_{\text{leak}}$ shift and/or $G_{\text{leak}}$ change | altered $Cl^-$ contribution to background clamp |
| CCCP | $G_{\text{leak}}$↓ (weaker clamp) and/or $\tau_{\text{rec}}$ ↑ | PMF collapse reduces active buffering; slows recovery |

```
MATLAB CODE

% run_MTP2_pumpleak.m
% Pump–leak model with concentration-dependent MTP2 bias and 10 s light pulse.
% Save this file, then run:  run_MTP2_pumpleak

clear; clc; close all;

%% Time axis (match your experiments: light at 120 s for 10 s)
t0     = 0;
t_end  = 380;
t_on   = 120;
t_off  = 130;   % 10 s pulse
tspan  = [t0 t_end];

%% Core electrical parameters (order-of-magnitude; tune as needed)
p.C      = 1.10e-13;    % F (example: B. subtilis total capacitance)
p.Gleak  = p.C/10;      % S -> sets fast tau = C/G ~ 10 s
p.Eleak  = -70e-3;      % V baseline set-point (adjust per condition)

% Slow recovery time constant (captures minutes-scale return)
```

```
p.tau_rec = 180;        % s (tune: 150–400 s)

% Light timing
p.t_on  = t_on;
p.t_off = t_off;

%% Concentrations to simulate (ug/mL)
c_list = [0 2 5 10];

% Define the concentration dependence of the MTP2 bias current.
% Here: linear in c with two amplitudes (dark vs light).
% You can replace with a Hill/saturating function if needed.
p.c_ref = 10;            % ug/mL reference
p.Idark_ref  = -2.0e-14; % A at 10 ug/mL (NEG -> hyperpolarizing bias)
p.Ilight_ref = +1.2e-14; % A at 10 ug/mL during light (depolarizing bias)

%% Initial condition (start near Eleak or your measured baseline)
V0 = -70e-3;  % V

%% Solve and plot
figure; hold on;
for k = 1:numel(c_list)
    c = c_list(k);
    p.c = c;

    opts = odeset('RelTol',1e-6,'AbsTol',1e-9,'MaxStep',1.0);
    [t, V] = ode45(@(t,V) rhs_pumpleak(t,V,p), tspan, V0, opts);

    plot(t, 1e3*V, 'LineWidth', 2); % mV
end

% Mark light pulse
yl = ylim;
patch([t_on t_off t_off t_on],[yl(1) yl(1) yl(2) yl(2)], ...
      [1 1 0], 'FaceAlpha', 0.15, 'EdgeColor', 'none');
uistack(findobj(gca,'Type','patch'),'bottom');

xlabel('Time (s)');
ylabel('V_m (mV)');
legend('0','2 \mug/mL','5 \mug/mL','10 \mug/mL','Location','best');
title('Pump–leak + MTP2 bias (10 s light pulse)');
```

```
grid on; box on;

%% ---- Local function: ODE RHS ----
function dVdt = rhs_pumpleak(t, V, p)
% Implements: C dV/dt = -Gleak (V - Eleak) + I_MTP2(t,c)

    % --- Concentration scaling (linear) ---
    scale = (p.c / p.c_ref);

    Idark  = p.Idark_ref  * scale;
    Ilight = p.Ilight_ref * scale;

    % --- Light gating (10 s pulse) ---
    is_light = (t >= p.t_on) && (t <= p.t_off);

    % --- Slow recovery of the bias after the light pulse ---
    % During light: apply Ilight
    % After light: relax exponentially from Ilight back to Idark with tau_rec
    if is_light
        Imtp2 = Ilight;
    elseif t > p.t_off
        Imtp2 = Idark + (Ilight - Idark)*exp(-(t - p.t_off)/p.tau_rec);
    else
        Imtp2 = Idark;
    end

    % --- Pump-leak dynamics ---
    dVdt = (-p.Gleak*(V - p.Eleak) + Imtp2) / p.C;

    % Safety: avoid NaN/Inf propagation
    if ~isfinite(dVdt)
        dVdt = 0;
    end
end
```

#### S4. Supplementary Kanamycin susceptibility data

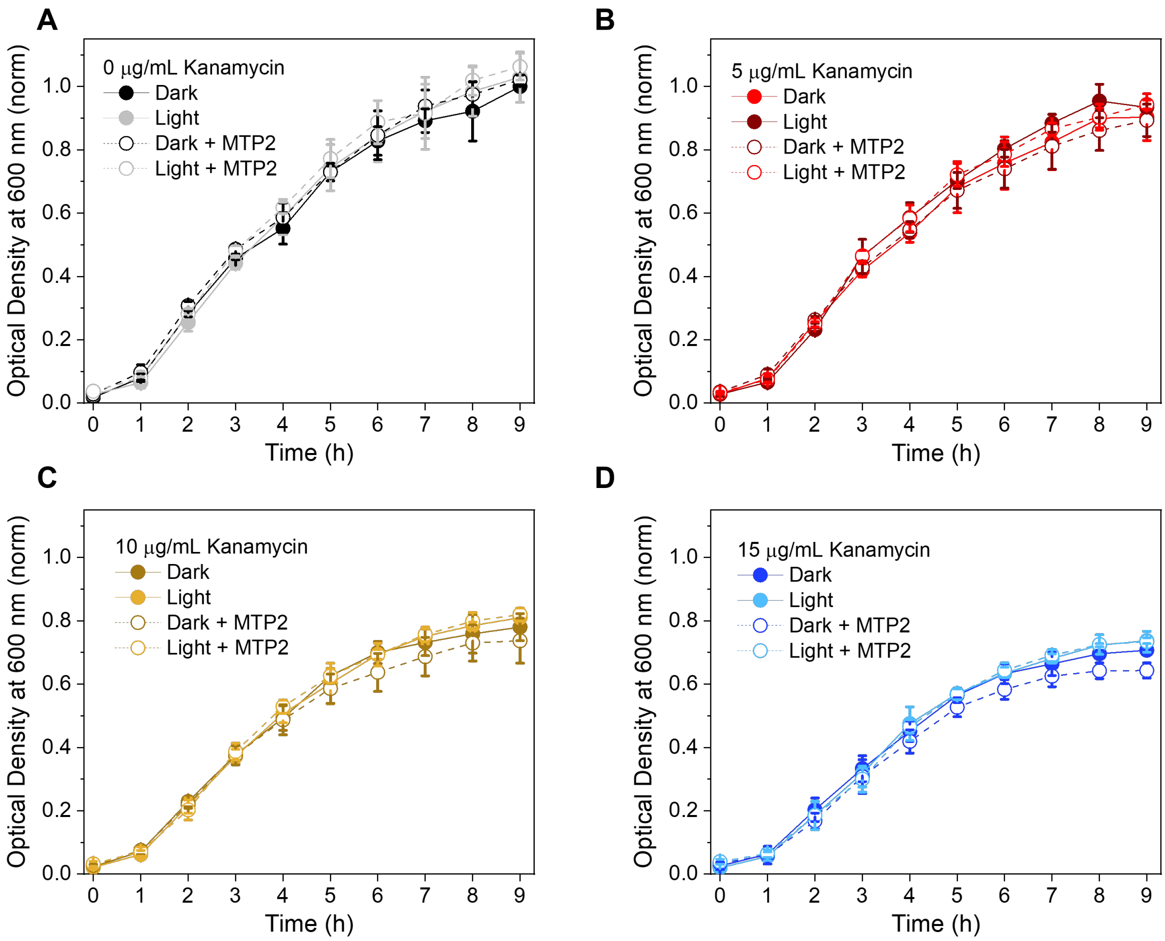


***Figure S5****: Growth curves of B. subtilis cells as monitored by the optical density at 600 nm, with and without 5 µg/mL MTP2, and with and without intermittent illumination at 470 nm, at kanamycin concentrations (A) 0 µg/mL, (B) 5 µg/mL, (C) 10 µg/mL, and (D) 15 µg/mL. These growth curves were normalized relative to the average final optical density at 0 µg/mL kanamycin under dark conditions.*

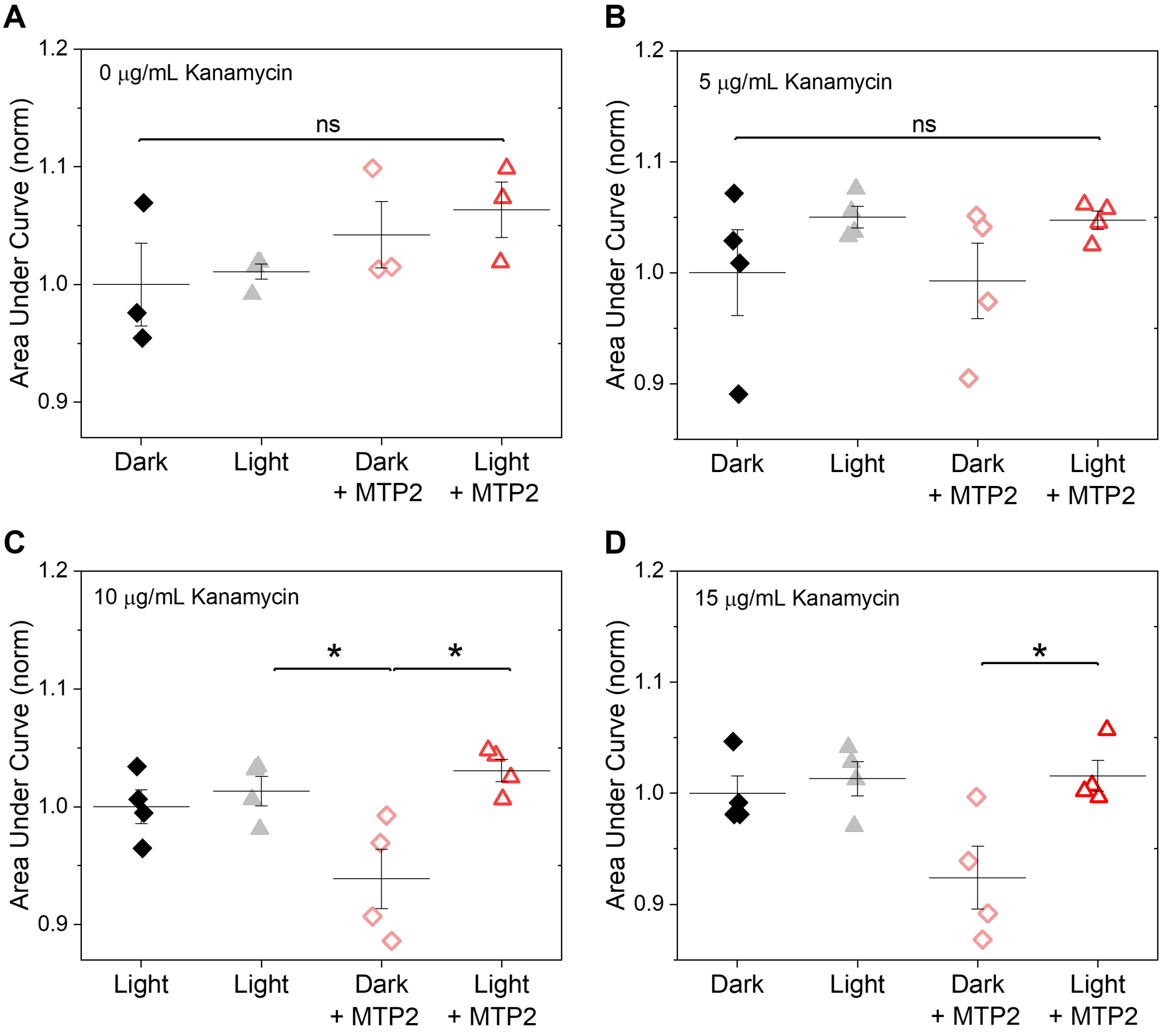


***Figure S6****: Areas under the growth curves of B. subtilis cells as monitored by the optical density at 600 nm (as seen in Figure 6A of the main text and Figure S5), for each condition with and without MTP2, and with and without intermittent illumination at 470 nm, at kanamycin concentrations (A) 0 µg/mL, (B) 5 µg/mL, (C) 10 µg/mL, and (D) 15 µg/mL. These data points were each normalized relative to the average area under the curve under dark conditions without MTP2 for each kanamycin concentration. For data sets with normally distributed points (A, C), the results of ANOVA statistical significance tests are shown. For data sets with non-normally distributed points (B, D), the results of Mann-Whitney U tests are shown.*
